# PATTIE: Publication access through tiered interaction & exploration

**DOI:** 10.1101/807404

**Authors:** Michael Segundo Ortiz, Mengqian Wang, Kazuhiro Seki, Heejun Kim, Javed Mostafa

## Abstract

In this work we present Publication Access Through Tiered Interaction & Exploration (PATTIE) – an information foraging, sense-making, and exploratory spatial-semantic information retrieval (IR) system (http://pattie.unc.edu/plos). Non-spatial, spatial IR systems, and some recent studies focused on their principal functions are discussed and compared. To interactively work through a use-case from the biomedical domain, instructions are provided for readers to conduct exploratory searches directly on the PLOS archive based on the software embedded in the online version of this paper (http://vzlib.unc.edu/software/). To carefully evaluate some of the critical parameters of the PATTIE algorithm, and the core functions of the implemented system, a set of experiments were conducted. Along with details on the experimental methods and their rationale, key findings from the experiments are analyzed and presented. Finally, with an eye toward the future of software-embedded scientific papers, their potential benefits for supporting direct engagement with scientific content, replication, and validation are discussed.

## Introduction

Information retrieval (IR) systems are essential tools for finding relevant documents. Current IR systems dominantly adopt the ranking-based retrieval model, which returns a list of documents ranked in descending order of predicted relevance for a user query (i.e., search keywords). Such an architecture and information access point rely on the user having and understanding on how and what to search for. The evidence base suggests that this is often an incorrect assumption to make [1, 2]. Despite a user overcoming these assumptions by searching appropriately, a Search Engine Result Page (SERP) can retrieve an extraordinarily large set of documents which makes it difficult to locate and comprehend all the relevant information. This is especially true for the biomedical domain given the ever-growing body of the literature; a position that IR researchers have been discussing for decades now.

For instance, consider a scenario where a researcher unfamiliar with the PLOS digital archive is curious to understand the topical structure. Without issuing a query, a dynamic spatial-semantic table-of-contents can be generated as shown in Fig 1. The PATTIE Map presents the most recently published topical content within the PLOS digital archive. We will demonstrate the access point, mechanism of navigation, and information acquisition in the *System design and implementation details* and *Discussion* sections.

**Fig 1.**
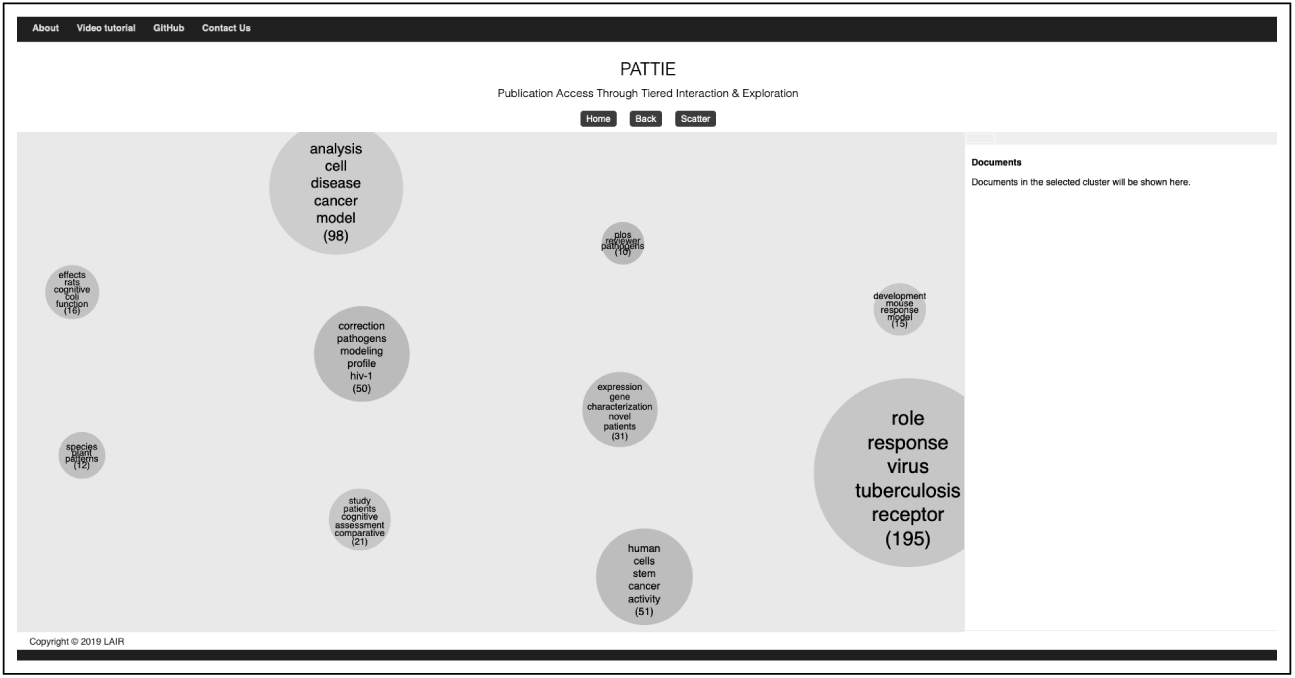
A screenshot of PLOS digital archive PATTIE Map, no query issued.

Alternative modes of access or metaphors for representing and presenting information spaces in IR that incorporate spatial-semantic context may prove beneficial, and have been discussed with works on exploratory search, sense-making, information foraging [3–9], and visualization of concept spaces [10]. Moreover, when we consider individual differences in verbal and spatial reasoning abilities, direct manipulation of sequential, 2D, and 3D interfaces, retrieval performance, and user satisfaction, it becomes apparent that the sequential list of relevant documents for a state-of-the-art IR system is no longer the state-of-the-art in cases where users are *not engaged* in look-up or transactional retrieval [11–22]. Such kind of information-seeking behavior, and systems to support it, do not require a specific query and typically have some mechanism to substantiate user intent by, for example, providing spatially-encoded keywords, categories, or clusters. Therefore, users can interact with what they believe to be pertinent instead of formulating a potentially ambiguous or improperly scoped query that results in a *filter bubble*. For an in-depth review on work related to these mechanisms and the differences between information visualization and information navigation enabled by visualization, please see [23].

Researchers intimately understand the rapidly accelerating growth and diversity of scholarly content, and according to Heap’s Law [24], as more research text is gathered, the discovery of the full vocabulary becomes insurmountable and thus proper query formulation becomes somewhat of a linguistical *arm’s race* as we will see in a use-case involving a researcher developing an outline for a review article in the *Discussion* and *Use-Case* section.

This paper tackles these problems and introduces a dynamic cluster-based browsing system adopting the Scatter/Gather paradigm [25–27], named *Publication Access Through Tiered Interaction & Exploration* (PATTIE). Scatter/Gather was developed with *information foraging theory* in mind [8, 9]. The theory sought to mathematically formalize the trade-offs between information acquisition and cognitive load. Thus, by topically organizing information, cognitive load could theoretically be reduced while maximizing information gain.

With/without an initial query, PATTIE dynamically generates topical clusters on the fly and visualizes the results for intuitive navigation via a spatial-semantic table-of-contents metaphor for scholarly digital archives. In the remaining sections, we will describe system architecture, evaluation of the unsupervised machine learning pipeline, and a use-case demonstrating the power of PATTIE.

## System design and implementation details

Our dynamic cluster-based document browsing system, PATTIE, for the PLOS archive was built on our prototype [23]. The main architecture is retained but the interface was given a few updates for better presentation and usability, including a function to maintain user’s selection of clusters throughout a session, which is critical in terms of accessibility and for users who may have low spatial memory [12, 14, 28]. For completeness, the following describes not only the differences from the prototype but the complete details of design and implementation of PATTIE.

### Architecture

Fig 2 depicts PATTIE’s design. The server side system is implemented by the Flask web framework [29]. The document collection (PLOS) is directly served through PLOS API [30], which eliminates the need to update the database at the server side and assures that the users can explore the most updated data. The client side is built on a JavaScript visualization library D3.js [31]. Ajax is implemented on the client side to asynchronously communicate with the server while content is dynamically explored, maintaining user work space without reloading the web page.

**Fig 2.**
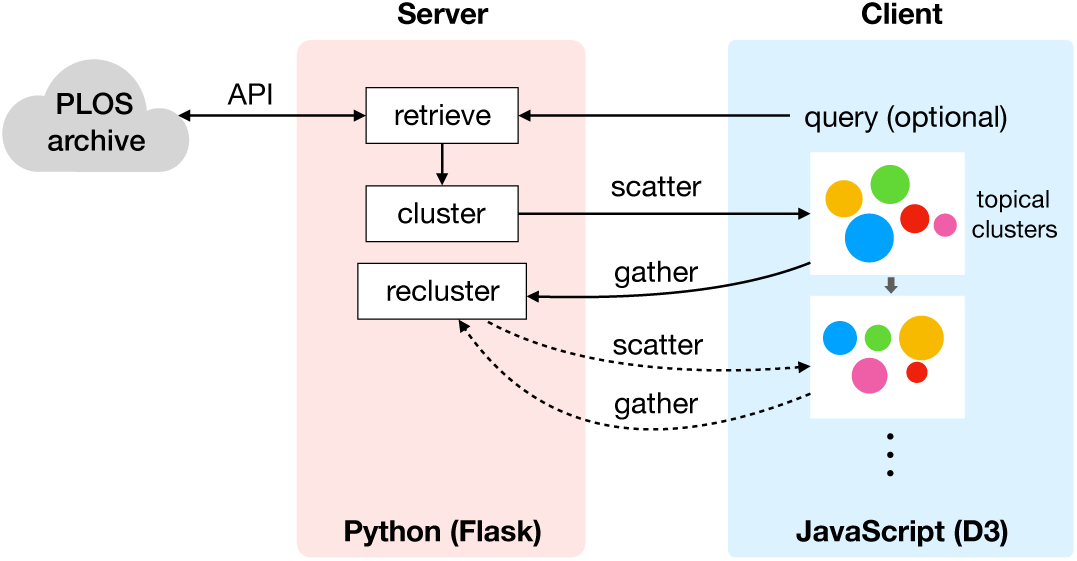
Design of PATTIE architecture.

### Server-side: Data Retrieval and Initial Clustering

As with the standard of the modern search systems, PATTIE presents a text box for a user to type in a query (Fig 3) although it can also initiate a process of information exploration *without* a query. When a search terms are provided, PATTIE retrieves *N* latest articles that map to the search terms. in any textual fields including titles, abstracts, and body texts. When no search terms are provided, PATTIE retrieves *N* latest articles indexed in the PLOS archive in order to provide the user a mechanism for archive sense-making. *N* is fixed to a constant in order to dynamically cluster archival content in constant time which facilitates real-time processing.

**Fig 3.**
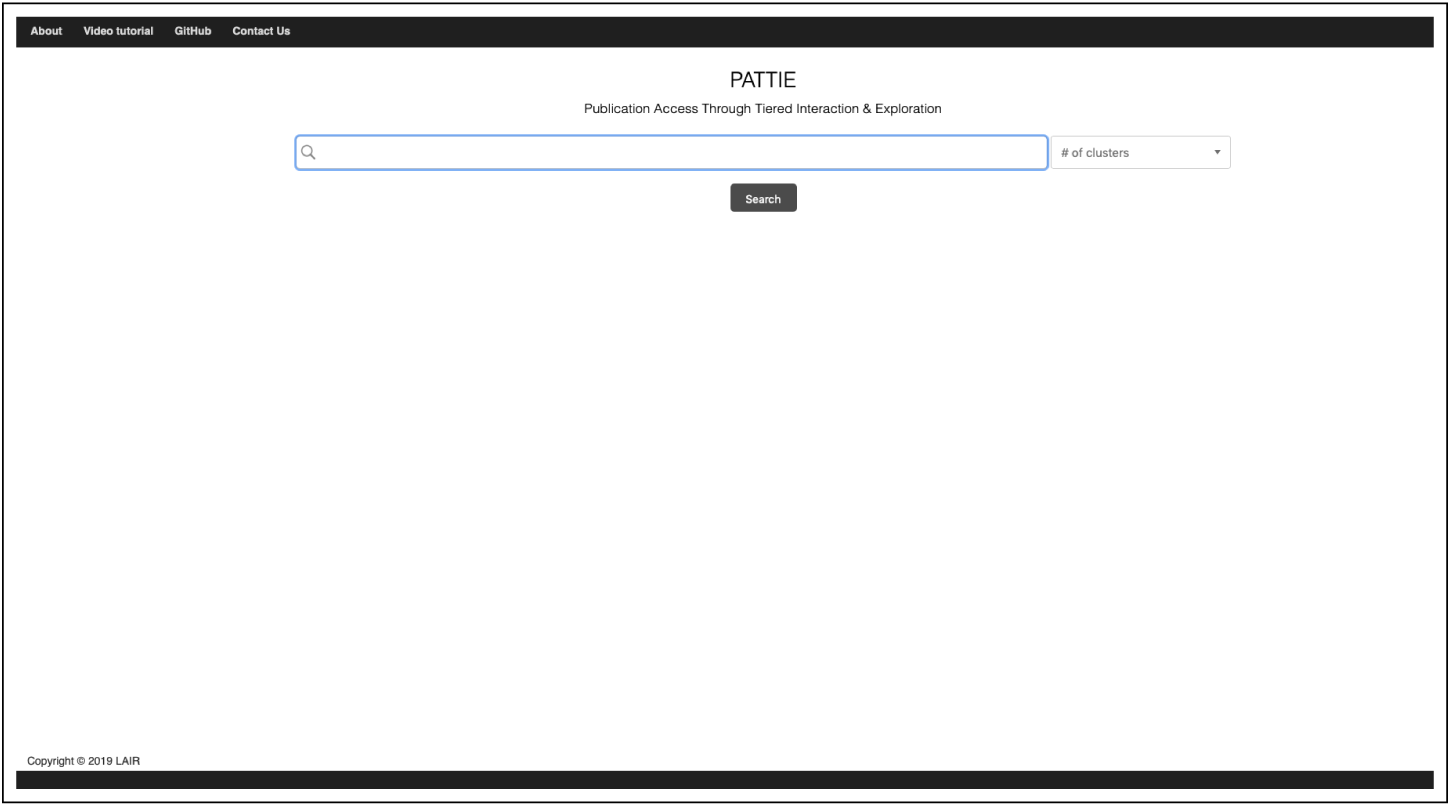
Screenshot of the PATTIE search page.

To some extent, this is similar to the idea of mini-batch *k*-means [32] which has been observed to perform significantly faster than *k*-means while still converging on a similar clustering solution. Instead of complete randomness, however, PATTIE focuses on index recency as biomedical researchers are generally concerned with emerging concepts in their field. Limiting the number of documents by *N* may have an impact on the the resulting cluster structure and its quality. However, we assume that the effect is limited when *N* is set to a sufficiently large value. We will empirically investigate the validity of this assumption in the *Discussion* section.

As for query language, PATTIE accepts a wide range of query syntax understood by the underlying PLOS API. Currently, the system retrieves concatenated titles and abstracts, but other indexed fields are available and we plan to study their use in future work.

After mapping the query to the archive and retrieving a document set, PATTIE executes a unsupervised machine learning pipeline that is in sequential order below. The pipeline was evaluated and it was concluded that PATTIE can partition information spaces into coherent clusters [33]. For a more detailed description of the pipeline, please see [23].

1. Keyword discovery: The system first analyzes prominent terms or features for document representation via Vector Space Modeling (VSM) and statistical term weighting to generate a matrix *M*.
2. Latent semantic analysis (LSA) [34]: The previous keyword discovery step decreases the vocabulary size and thus the dimensionality of document vectors. In order to further reduce the dimensionality and to discover latent associations among keywords, LSA is applied to *M*.
3. Clustering: logical partitions are predicted by the *k*-means++ algorithm [32].

Then, PATTIE generates a set of keywords to describe each cluster for the next visualization stage as follows. First, the centroid of each cluster in the LSA-reduced space is transformed back to the original term-document space, which can be thought of as a pseudo document vector. From the vector, a fixed number of keywords with values meeting the threshold are selected as cluster labels.

In addition, the centroids in the LSA-reduced space are transformed to 2D coordinate space by t-Distributed Stochastic Neighbor Embedding (t-SNE) [35] for presentation. Only the cluster membership, cluster descriptors (keyword set), 2D coordinates of the clusters, and basic bibliographic information (author names, article titles, publication dates, and journal titles) of the search results are sent to the client to keep the data traffic minimal, and other information, such as the term-document matrix *M*, is retained as session data on the server.

### Client-side: Visualization

To create an intuitive, user-friendly interface, we rely on Shneiderman’s mantra [36] for visual information seeking—*overview first, zoom and filter, then details on demand*—while designing the PATTIE interface. According to the mantra, the PATTIE system provides overviews of clusters first and shows details according to users’ interests. There are two panels and buttons for Scatter/Gather (Fig 4) on the interface. The left panel (“PATTIE Map”) displays the partitioned information space and the right panel (“document panel”) displays the corresponding documents. Color encoding is used for clusters as well as the document panel to provide a cue for more efficient navigation.

**Fig 4.**
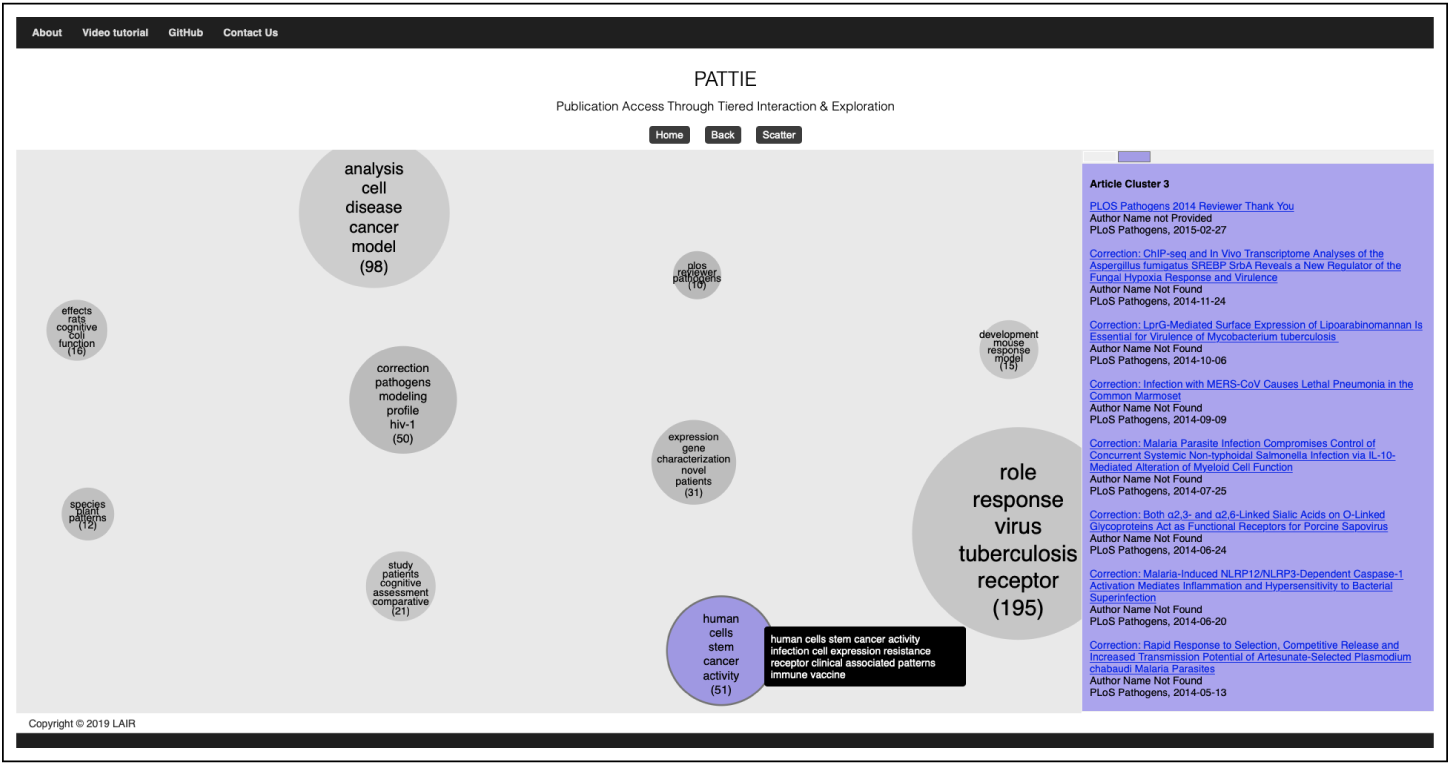
PATTIE Map with selected cluster within the PLOS digital archive, no query issued.

For effective Scatter/Gather visualization, it is crucial how to place and present clusters on the PATTIE Map, as it can have a huge influence on how users perceive and interpret clusters. Thus, we locate the cluster centroids based on virtual coordinates in semantic space constructed by t-SNE, which reflects their relative semantic relationships. In other words, clusters that are closer to each other are semantically more similar than clusters that are farther apart. In addition, in order to expose the sense of spatial encoding to users, clusters are first placed in the center of the PATTIE Map and immediately “scattered” to their coordinates.

In the PATTIE Map, a circle represents a cluster, and the area of the cluster is proportional to the number of documents that belong to the cluster. The color of a cluster also indicates the virtual semantic coordinates of the cluster. Clusters are initially shown in gray but their color changes to the one determined by the coordinates of their centroid when clicked. Specifically, we used the HSV (hue, saturation, and value) color model and considered the center of the PATTIE Map as the origin. Hue and saturation are determined by the angle and the *ℓ*^2^-norm of the vector from the origin to the centroid of a corresponding cluster, whereas value is fixed to a constant.

Five representative keywords (cluster descriptors) are presented within each cluster along with the number of documents belonging to that cluster in parentheses. When a user hovers the mouse pointer over a cluster, 15 descriptors including the five are shown as a tooltip to help him/her assess the relevance of the cluster. Also, the corresponding bibliographies with a hyperlink to the original article registered in the PLOS digital archive temporarily appear in the document panel as a user hovers their mouse over the cluster. If a user clicks a cluster(s) of interest (Gather), the bibliographies remain in the document panel until the cluster is deselected. If a user selects multiple clusters, the cluster tabs located at the top of the document panel help the user navigate the documents belonging to each cluster. The cluster on the PATTIE Map and its corresponding tab in the document panel has the same color to facilitate intuitive navigation.

After a user decides which clusters to choose for the re-clustering using the above features, he/she can start the re-clustering by clicking the “Scatter” button. The chosen clusters are first moved to the center of the PATTIE Map shown as an animation, computationally re-clustered on the server-side as described in the next section, and then scattered to the coordinates of new clusters. This animation should be able to help users conceptualize the Scatter/Gather process, although they are new to the idea. Users can iterate and/or restart this Scatter/Gather process until their information needs are satisfied.

The user can also choose to go back to the previous state by clicking the “Back” button. When clicked, the scattered clusters move back to the center of the PATTIE Map and then the previously presented clusters are scattered again, where the previously chosen clusters are kept selected and are shown in the document panel. In other words, their previous “mental map” is conserved so that the user can focus on exploration and comprehension of the information space instead of re-selecting clusters.

### Server-side: Re-clustering

After PATTIE receives a set of selected clusters, the system retrieves the IDs of documents belonging to the clusters from the session data stored on the server and carries out the unsupervised machine learning pipeline previously described above. These processes may appear redundant and unnecessary. However, we must emphasize that PATTIE relies not on an information visualization per se, but an information visualization that enables iterative navigation. We believe such a mechanism affords user(s) to comprehend research concepts, logical connections, relevance, and scope, as the user(s) narrows down toward a latent information target.

This mechanism is crucial for PATTIE to identify a *new* set of keywords, which would be more narrowly focused from the previously identified keywords and, consequently, yielding more scoped sub-clusters. Sub-cluster information is then computed via the pipeline and sent to the client side for visualization and iterative navigation.

## Discussion

### Sampling-based Clustering

For cluster-based document browsing, such as Scatter/Gather, it is vital that clustering is completed in real time irrespective of the size of the archive or search result. There are existing approaches running in a constant time [26, 37], which however rely on a pre-computed, static hierarchy of categories. Dynamicity is the core of PATTIE and the result of re-clustering should dynamically change upon users’ selection of clusters or the underlying document collection. That is, as the PLOS archive evolves, so too should the PATTIE Map, effectively providing users an evolving spatial-semantic table-of-content metaphor for the life of the archive.

We proposed a simple strategy to retrieve only the *N* latest articles for a given query such that clustering completes approximately in a constant time for fixed *N* [23]. The relation between the size of *N* and the quality of clusters was informally studied in order to find the value of *N* which could produce as good clusters as those produced from the entire search results. Here, we repeated the same experiment to reexamine the relation more rigorously with significance tests. Also, we compared standard *k*-means and mini-batch *k*-means to examine if further speed-up could be achieved. For these experiments, we use the same data set and an evaluation criterion (Adjusted Mutual Information (AMI) [38]) as the previous work [23].

#### Results

Fig 5 displays the relationship between the number of sampled (latest) documents *N* and cluster quality, while *N* was gradually increased from 200 to 12,530. For each *N*, we repeated clustering for 20 times and plotted the mean AMI with the standard deviation as an error bar. In the figure, “Title”, “Abstract”, and “Full text” indicate the results produced by titles, titles and abstracts, and titles and abstracts and body text, respectively. The observations are similar to what was reported before [23]; AMI sharply improved as the sample size increased up to 2,000 for Title and Abstract and then stabilized while full-text data was not as effective.

**Fig 5.**
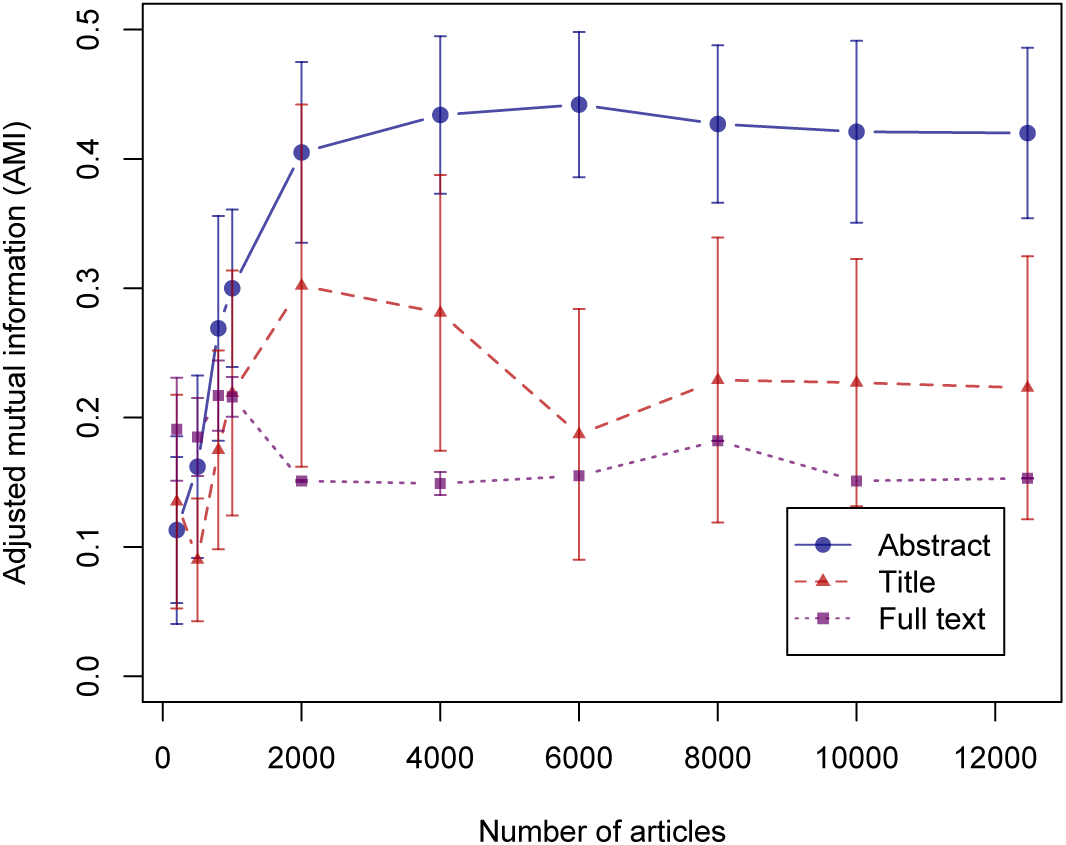
Cluster quality against increasing dataset size and type of data.

The result indicates that more information does not necessarily translate to greater performance in clustering documents potentially due to more irrelevant words brought in with full text. A difference from our previous work [23] is that using titles only did not yield as good clusters as using abstracts. In fact, the difference between Abstract and Title was found statistically significant at the significance level of 0.01 for *N ≥* 500 by Welch’s unequal variances *t*-test.

We also examined the processing time for Title, Abstract, and Full text in Fig 6, which was measured as the total time required for constructing a tf-idf matrix, applying VCGS and SVD, and clustering excluding the time to load data into memory. Using Full text has a clear disadvantage in processing time as well as cluster quality. Based on these observations, our current system retrieves 500 latest articles (i.e., *N* = 500) and uses titles and abstracts for discovering topical clusters.

**Fig 6.**
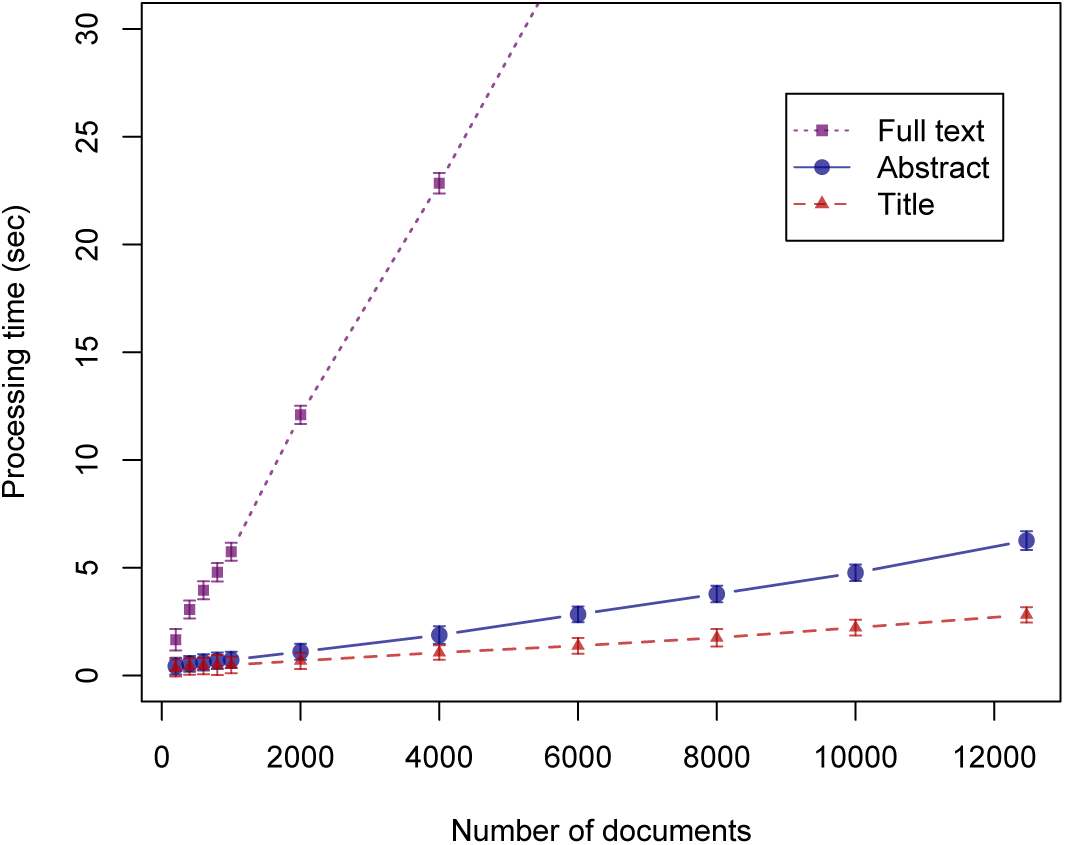
Relation between the number of documents and processing time.

Here, it should be noted that the above experiment only examined cluster quality, not the accessibility of information. That is, if relevant articles were not among the latest *N* articles, one can never find the relevant articles. Also, while we chose to retrieve *N* latest articles, one could use *N* random articles instead, which may work better depending on user information need. To avoid such issues, *N* should be ideally equal to the size of the document collection or search result, which is however difficult to process in real time depending on the data size as observed in Fig 6. *N* is currently limited to a manageable size balancing cluster quality and processing time, but we plan to increase it by employing more efficient data structure and distributed processing in future work.

#### Comparison with mini-batch *k*-means

Mini-batch *k*-means [32] uses a mini-batch optimization for *k*-means clustering, which has been reported to greatly reduce computation time but still achieve a solution close to the standard *k*-means algorithm. The algorithm first takes *b* random samples as a mini-batch and each sample in the mini-batch is assigned with the nearest centroid and then each centroid is updated per-sample basis. The assignment and update steps are repeated for predetermined times or until convergence.

As a real-time spatial-semantic system, it is vital for PATTIE to have a minimal turnaround time with a balance between accuracy and latency in mind. To this end, we investigated mini-batch *k*-means as an alternative. Specifically, we performed the same experiment as the previous section changing the number of sample documents *N* to measure the clustering performance in AMI using mini-batch *k*-means in order to compare it to the standard *k*-means. Only titles and abstracts were used for this experiment. Figs 7 and 8 show the results.

**Fig 7.**
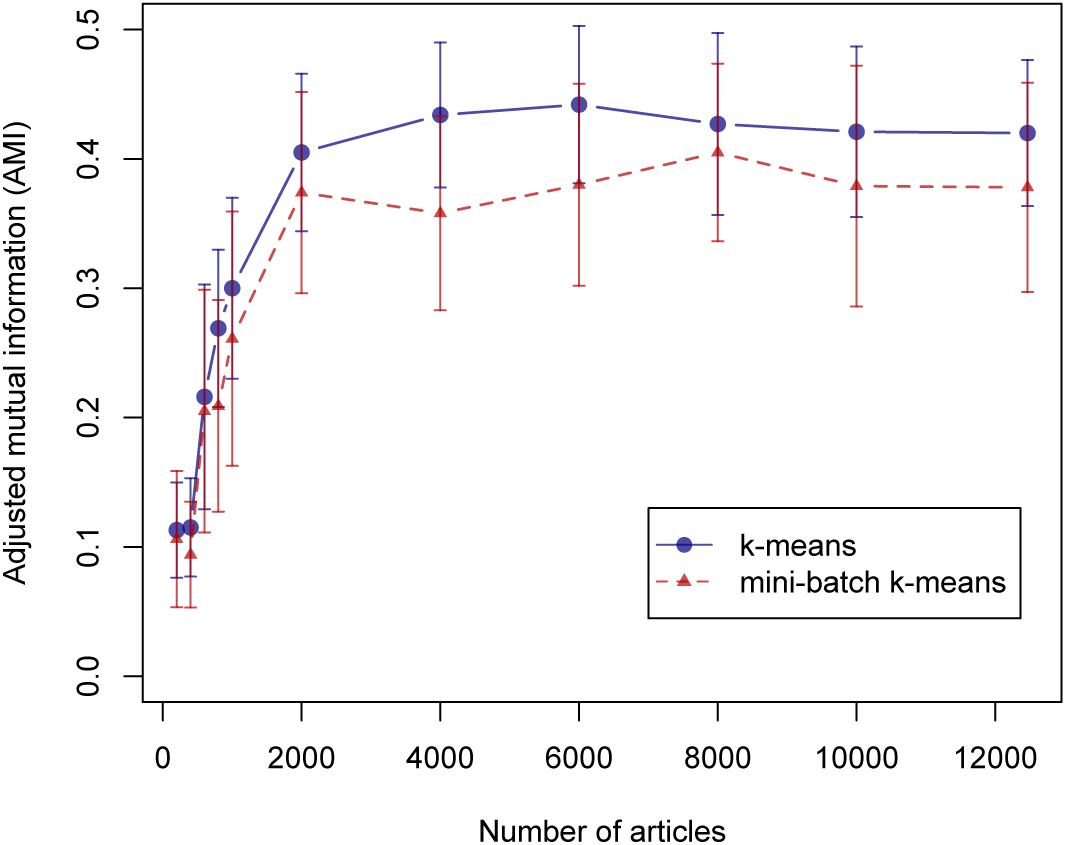
Relation between the number of documents and processing time.

**Fig 8.**
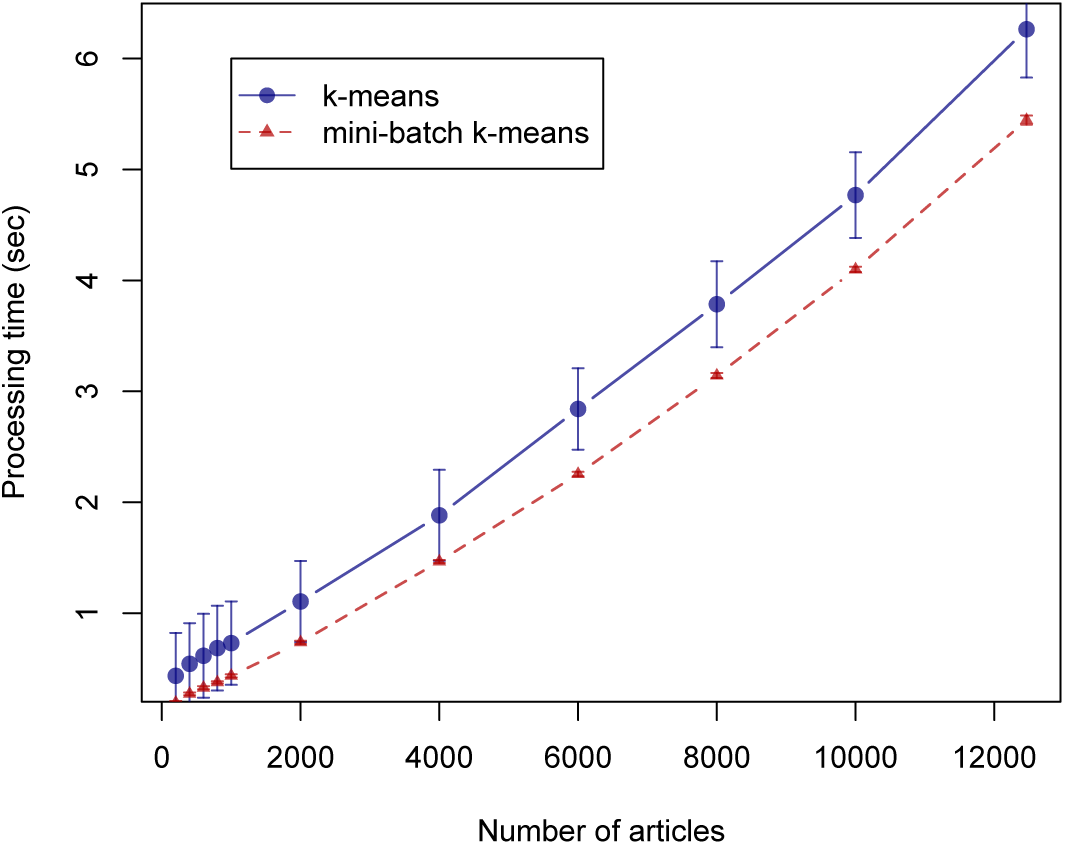
Relation between the number of documents and processing time.

Although *k*-means achieves slightly higher AMI, the difference was found to be not statistically significant and mini-batch *k*-means runs slightly faster (approximately 0.5 seconds) irrespective of *N*. Their processing times did not differ greater because a large portion of the processing time (64∼90% for *N* = 2, 000) is accounted for constructing a tf-idf matrix and applying VCGS and SVD.

Overall, mini-batch *k*-means algorithm brings a slight increase in speed with insignificant difference in clustering performance. While it is a valid alternative, more work needs to be done on the processes including tf-idf matrix construction for further improvement. Therefore, our current system adopts the standard *k*-means.

### Use Case

In the following, we will demonstrate how PATTIE can be used to explore the PLOS digital archive with respect to a use-case involving a university student’s workflow for creating an outline for a review article on *the latest advancements in modulating CRISPR guided gene editing* by using a PATTIE Map for navigating and foraging on the PLOS digital archive.

There are a few essential steps required for the student to accomplish the task of creating an outline for a review article on *the latest advancements in modulating CRISPR guided gene editing*. The following items are logically ordered thought processes of the student while anticipating, and engaging in, this information-seeking task.

1. What search terms do I use?
2. Am I being comprehensive enough?
3. Should I re-formulate my search terms to make sure?
4. I think I have covered all the potential queries.
5. How many articles have I gathered?
6. Which articles are precisely relevant to my research question?
7. I am going to focus only on this subset as they are pertinent.
8. How best can I logically organize the outline in terms of the biological concepts that are thematic?

The student ponders on the subject matter and realizes that the potential vocabulary needed for proper query formulation will take much time and likely exceeds their level of current knowledge according to Heap’s Law [24]. However, upon issuing the simplistic query “*crispr*” to PATTIE, much of this uncertainty is mitigated by allowing PATTIE’s unsupervised machine learning pipeline to analyze the vocabulary and generate a map as shown in Fig 9

**Fig 9.**
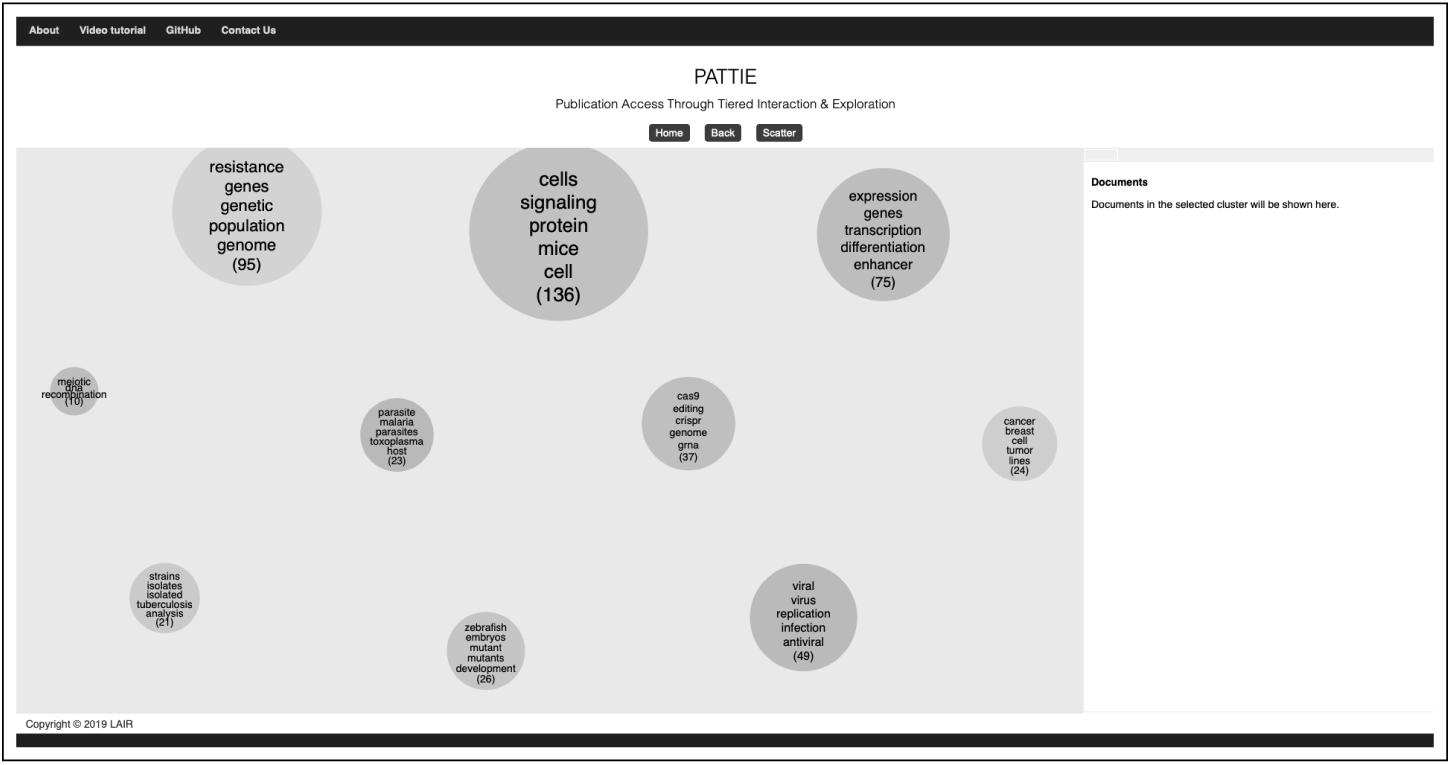
CRISPR PATTIE Map within the PLOS digital archive.

A PATTIE Map eliminates much of the cognitive load associated with the workflow above. To access the information space, only a simple query is needed. After reflecting for a moment on the information space, the student is cued in on the keywords *Cas9, editing, crispr, genome* and *grna* as shown in Fig 10. At a very high-level, the student understands that *CRISPR associated protein 9 (Cas9)* is the essential mechanism for cutting DNA, and is more precisely an endonuclease enzyme [39]. Moreover, *editing, crispr* and *genome* are of course self-explanatory with respect to the original research question regarding modulation of CRISPR. Lastly, the student recalls that *Guide RNAs (grna)* are responsible for the insertion and deletion of nucleotide bases associated with the engineering involved in CRISPR technology [40].

**Fig 10.**
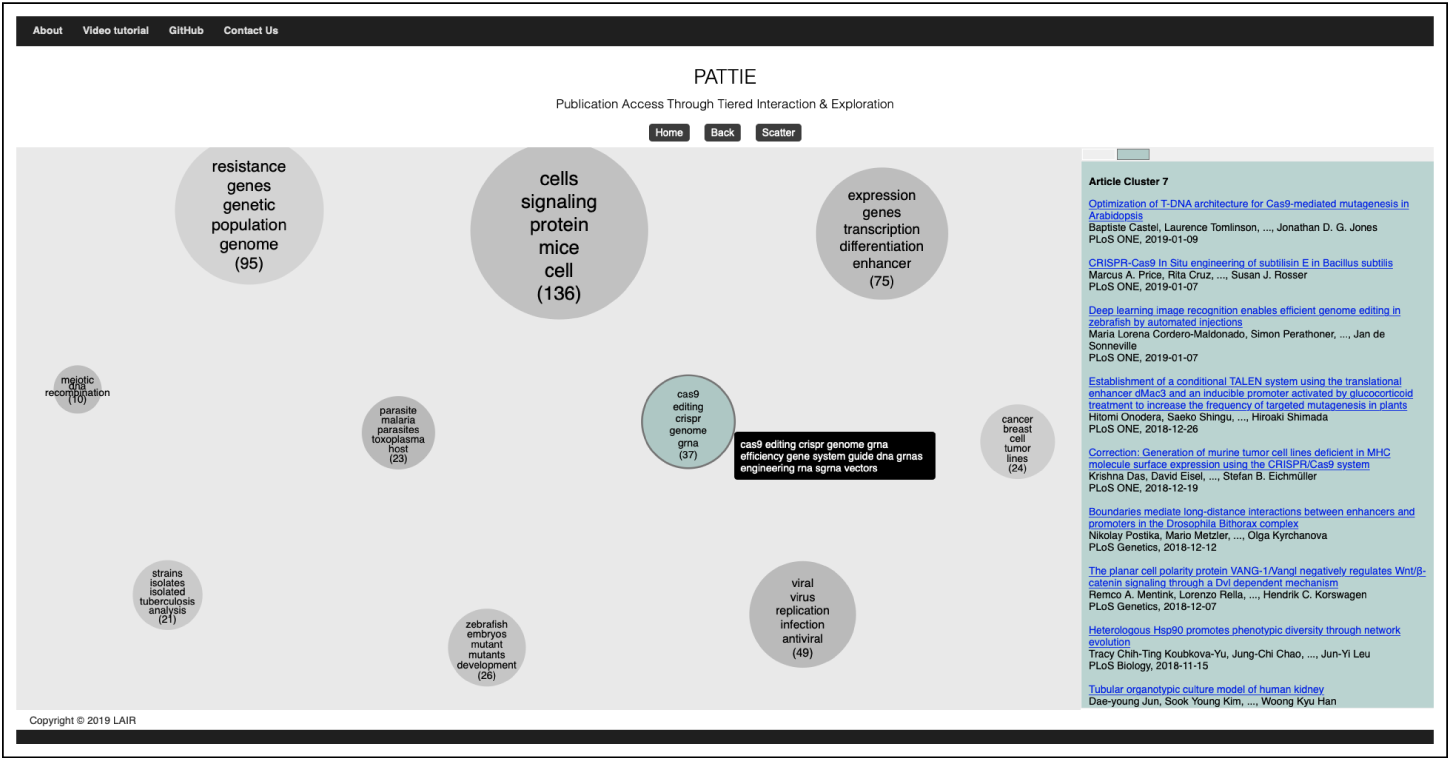
User-scoped PATTIE Map for the query *crispr* with cluster selection color encoded.

The student hovers their mouse over the cluster and is even more certain on the selection with respect to the *details on demand* keywords that include *efficiency* and *engineering*. A *scatter* phase is initiated for further inspection. The student examines the more scoped information space and begins selecting clusters with the following thought processes itemized below, and as shown in Fig 11:

**Fig 11.**
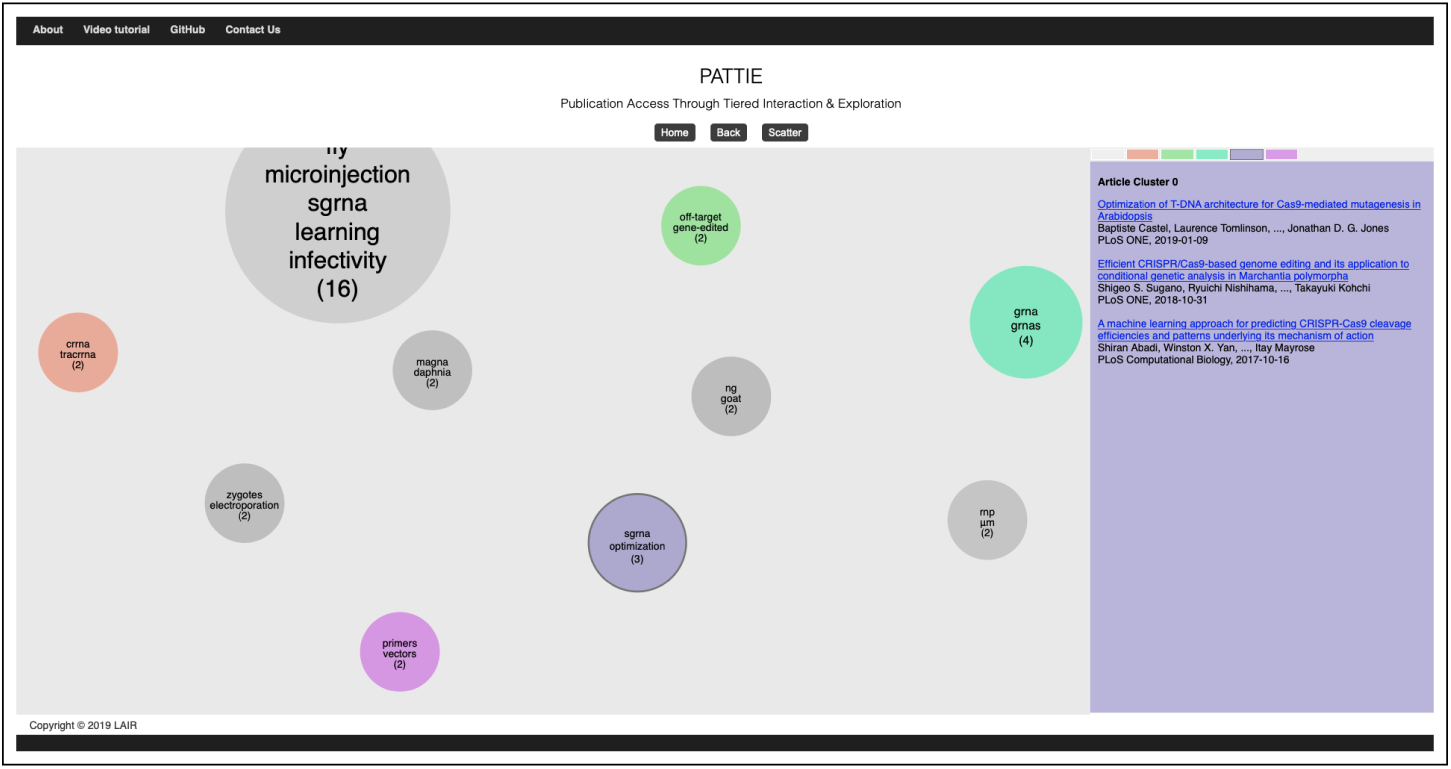
Student narrowing the information targets and reasoning through potential connections for developing the outline.

- *crrna, tracrrna* – CRISPR Associated RNA (crrna) and Tran-activating CRISPR associated RNA (tracrrna) form complexes for defense and immunity within bacteria. This may be useful for my outline in understanding how innate biological modulation is being studied for CRISPR-guided therapy and its modulation in synthetic systems.
- *primers, vectors* – Primers and vectors are used to amplify genomic sequences that are then delivered to the molecular target via a bacterial vector. If these are being engineered in novel ways then I would like to understand what the implications are in terms of modulating CRISPR-guided therapy.
- *sgrna, optimization* – As I recall, insertion and deletion of nucleotide bases is an essential function of Guide RNAs, also referred to as Single-Guide RNAs, which are synthetic complexes that incorporate both crRNA and TracrRNA. This area of research may be examining the optimization of synthetic design. This feels important for understanding modulation of CRISPR-guided therapy.
- *off-target, gene-edited* – If CRISPR-guided therapy can result in off-target effects with unintended gene-edits then these studies are crucial to understanding how modulation of CRISPR activity will need to be further investigated.

The student has now scoped the information space by iterating through *Scatter/Gather* phases while focused only on the concepts and their logical connections as opposed to search mechanics. In other words, the student shifts attention to information processing and retrieval instead of search and access points. Moreover, while interacting with PATTIE, focus naturally gravitates toward learning about the information space and not at all about search parameters, query re-formulations, page numbers, or number of articles, because PATTIE cues the student on spatial-semantics that serve as a dynamic table-of-contents metaphor which partitions the space automatically and intelligently. Although, it must be noted that previous evidence indicates that direct manipulation such as operating spatial systems with non-spatial input devices (computer mouse) can cause issues for some users who have no experience with such systems in the past. However, this same evidence-base also demonstrated that individuals with varying levels of spatial reasoning ability can acquire this direct manipulation skill rather quickly [13, 28]. Thus, as with all information systems, a learning phase would be beneficial for certain users to process PATTIE Maps as demonstrated in this use-case.

## Conclusion

Spatial-semantic information retrieval is not a new concept. However, the tools for navigation and exploration of higher-dimensional (2D & 3D) information retrieval systems are difficult to find and/or non-existent that support researchers in what we would refer to as *Research Support* or *Research Analytics*. The earliest example of such a system was developed by Zhang et al. [10]. Essentially the authors argued that digital archives would eventually outpace the rate of human processing ability. Therefore, navigation tools that were directly *hooked in* to a live archive would provide users the spatial-semantic table-of-contents metaphor. The system was demonstrated as an *interactive paper/executable article* where the content and organizational separation of theory, method, and findings, from the actual data, was now one electronic entity. This early work motivated the vision of *Documents and (as) Machines* [41]. As a humble extension of this no longer available early work, we have built PATTIE in the same tradition. We hope to provide a tool to support *Research Analytics* that will enable insightful exploration of the PLOS Digital Archive. For more information on the concept of an *executable article* see *S1 Appendix*.

## Supporting information

### S1 Appendix: PATTIE Executable Article

A demonstration of a *machine document* is available at http://vzlib.unc.edu/software/.

## References

1. White RW, Kules B, Drucker SM, et al. Supporting exploratory search, introduction, special issue, communications of the ACM. Communications of the ACM. 2006;49(4):36–39.

2. Wilson M, Russell A, Smith DA, et al. mSpace: improving information access to multimedia domains with multimodal exploratory search. Communications of the ACM. 2006;49(4):47–49.

3. Marchionini G. Exploratory Search: From Finding to Understanding. Commun ACM. 2006;49(4):41–46. doi:10.1145/1121949.1121979.

4. White RW, Roth RA. Exploratory search: Beyond the query-response paradigm. Synthesis lectures on information concepts, retrieval, and services. 2009;1(1):1–98.

5. White RW, Marchionini G, Muresan G. Evaluating exploratory search systems. Information Processing and Management. 2008;44(2):433.

6. Kules B, Capra R, Banta M, Sierra T. What do exploratory searchers look at in a faceted search interface? In: Proceedings of the 9th ACM/IEEE-CS joint conference on Digital libraries. ACM; 2009. p. 313–322.

7. Capra RG, Marchionini G. The relation browser tool for faceted exploratory search. In: Proceedings of the 8th ACM/IEEE-CS joint conference on Digital libraries. ACM; 2008. p. 420–420.

8. Pirolli P, Card S. Information foraging in information access environments. In: Chi. vol. 95; 1995. p. 51–58.

9. Pirolli P, Schank P, Hearst M, Diehl C. Scatter/gather browsing communicates the topic structure of a very large text collection. In: Proceedings of the SIGCHI conference on Human factors in computing systems. ACM; 1996. p. 213–220.

10. Zhang J, Mostafa J, Tripathy H. Information retrieval by semantic analysis and visualization of the concept space of D-Lib® Magazine. D-lib Magazine. 2002;8(10):1082–9873.

11. Benyon D, Höök K. Navigation in information spaces: Supporting the individual. In: Human-Computer Interaction INTERACT’97. Springer; 1997. p. 39–46.

12. Swan RC, Allan J. Aspect windows, 3-D visualizations, and indirect comparisons of information retrieval systems. MASSACHUSETTS UNIV AMHERST DEPT OF COMPUTER SCIENCE; 1997.

13. Sebrechts MM, Cugini JV, Laskowski SJ, Vasilakis J, Miller MS. Visualization of search results: a comparative evaluation of text, 2D, and 3D interfaces. In: Proceedings of the 22nd annual international ACM SIGIR conference on Research and development in information retrieval. ACM; 1999. p. 3–10.

14. Chen C. Individual differences in a spatial-semantic virtual environment. Journal of the American society for information science. 2000;51(6):529–542.

15. Westerman SJ, Cribbin T. Cognitive ability and information retrieval: When less is more. Virtual Reality. 2000;5(1):1–7.

16. Westerman SJ, Cribbin T. Mapping semantic information in virtual space: dimensions, variance and individual differences. International Journal of Human-Computer Studies. 2000;53(5):765–787.

17. Koshman S. Comparing usability between a visualization and text-based system for information retrieval. Journal of Documentation. 2004;60(5):565–580.

18. González-IbáñTez R, Proaño-Ríos V, Fuenzalida G, Martinez-Ramirez G. Effects of a visual representation of search engine results on performance, user experience and effort. Proceedings of the Association for Information Science and Technology. 2017;54(1):128–138.

19. Peltonen J, Belorustceva K, Ruotsalo T. Topic-relevance map: Visualization for improving search result comprehension. In: Proceedings of the 22nd International Conference on Intelligent User Interfaces. ACM; 2017. p. 611–622.

20. Ruotsalo T, Peltonen J, Eugster MJA, Głowacka D, Floréen P, Myllymäki P, et al. Interactive Intent Modeling for Exploratory Search. ACM Trans Inf Syst. 2018;36(4):44:1–44:46. doi:10.1145/3231593.

21. Klouche K, Ruotsalo T, Micallef L, Andolina S, Jacucci G. Visual re-ranking for multi-aspect information retrieval. In: Proceedings of the 2017 Conference on Conference Human Information Interaction and Retrieval. ACM; 2017. p. 57–66.

22. Schleußinger M, Henkel M. Evaluating a Visual Search Interface. LIS Scholarship Archive. 2018;.

23. Ortiz M, Kim H, Wang M, Seki K, Mostafa J. Dynamic Cluster-based Retrieval and Discovery for Biomedical Literature. In: Proceedings of the 10th ACM Conference on Bioinformatics, Computational Biology, and Health Informatics (ACM BCB); 2019. p. 390–396.

24. Heaps HS. Information retrieval, computational and theoretical aspects. Academic Press; 1978.

25. Cutting DR, Karger DR, Pedersen JO, Tukey JW. Scatter/Gather: A Cluster-based Approach to Browsing Large Document Collections. In: Proceedings of the 15th Annual International ACM SIGIR Conference on Research and Development in Information Retrieval. SIGIR’92. New York, NY, USA: ACM; 1992. p. 318–329. Available from: http://doi.acm.org/10.1145/133160.133214.

26. Cutting DR, Karger DR, Pedersen JO. Constant Interaction-time Scatter/Gather Browsing of Very Large Document Collections. In: Proceedings of the 16th Annual International ACM SIGIR Conference on Research and Development in Information Retrieval. SIGI ’93. New York, NY, USA: ACM; 1993. p. 126–134. Available from: http://doi.acm.org/10.1145/160688.160706.

27. Hearst MA, Pedersen JO. Reexamining the Cluster Hypothesis: Scatter/Gather on Retrieval Results. In: Proceedings of the 19th Annual International ACM SIGIR Conference on Research and Development in Information Retrieval. SIGIR’96. New York, NY, USA: ACM; 1996. p. 76–84. Available from: http://doi.acm.org/10.1145/243199.243216.

28. Billingsley PA. Navigation through hierarchical menu structures: does it help to have a map? In: Proceedings of the Human Factors Society Annual Meeting. vol. 26. SAGE Publications Sage CA: Los Angeles, CA; 1982. p. 103–107.

29. Flask; [cited 8 September 2019]. In: Pallets [Internet]. Available from: https://palletsprojects.com/p/flask/.

30. PLOS API; [cited 8 October 2019]. In: Public Library Of Science [Internet]. Available from: http://api.plos.org.

31. Bostock M. D3.js - Data-Driven Documents; [cited 8 September 2019]. Available from: https://d3js.org.

32. Sculley D. Web-scale k-means Clustering. In: Proceedings of the 19th International Conference on World Wide Web. WWW’10. New York, NY, USA: ACM; 2010. p. 1177–1178. Available from: http://doi.acm.org/10.1145/1772690.1772862.

33. Ortiz MS, Seki K, Mostafa J. Toward Exploratory Search in Biomedicine: Evaluating Document Clusters by MeSH as a Semantic Anchor. CoRR. 2018; 1812.02129.

34. Landauer TK, Foltz PW, Laham D. An introduction to latent semantic analysis. Discourse processes. 1998;25(2-3):259–284.

35. van der Maaten L, Hinton G. Visualizing Data using t-SNE. Journal of Machine Learning Research. 2008;9:2579–2605.

36. Shneiderman B. The Eyes Have It: A Task by Data Type Taxonomy for Information Visualizations. In: Proceedings of the 1996 IEEE Symposium on Visual Languages. VL’96. Washington, DC, USA: IEEE Computer Society; 1996. p. 336–. Available from: http://dl.acm.org/citation.cfm?id=832277.834354.

37. Ke W, Sugimoto CR, Mostafa J. Dynamicity vs. Effectiveness: Studying Online Clustering for Scatter/Gather. In: Proceedings of the 32Nd International ACM SIGIR Conference on Research and Development in Information Retrieval. SIGIR’09. New York, NY, USA: ACM; 2009. p. 19–26. Available from: http://doi.acm.org/10.1145/1571941.1571947.

38. Romano S, Vinh NX, Bailey J, Verspoor K. Adjusting for chance clustering comparison measures. The Journal of Machine Learning Research. 2016;17(1):4635–4666.

39. Genome Editing Primer; [cited 10 September 2019]. In: NIH-NLM Genetics Home Reference. Available from: https://ghr.nlm.nih.gov/primer/genomicresearch/genomeediting.

40. Addgene CRISPR Guide; [cited 10 September 2019]. In: Addgene, the non-profit plasmid repository. Available from: https://www.addgene.org/crispr/guide/.

41. Mostafa J. Documents and (as) machines. Journal of the Association for Information Science and Technology. 2018;69(1):3–5. doi:10.1002/asi.23993.

